# Mechanism of targeted killing of P. aeruginosa by pyocins SX1 and SX2

**DOI:** 10.1101/2022.10.27.514055

**Authors:** Jiraphan Premsuriya, Khedidja Mosbahi, Iva Atanaskovic, Colin Kleanthous, Daniel Walker

## Abstract

*Pseudomonas aeruginosa* is a common cause of serious hospital-acquired infections, the leading proven cause of mortality in people with cystic fibrosis and is associated with high levels of antimicrobial resistance. Pyocins are narrow spectrum protein antibiotics produced by *P. aeruginosa* that kill strains of the same species and have the potential to be developed as therapeutics targeting multi-drug resistant isolates. We have identified two novel pyocins designated SX1 and SX2. Pyocin SX1 is a metal-dependent DNase while pyocin SX2 kills cells through inhibition of protein synthesis. Mapping the uptake pathways of SX1 and SX2 shows these pyocins utilize a combination of the common polysaccharide antigen (CPA) and a previously uncharacterized TonB-dependent transporter (TBDT) PA0434 to traverse the outer membrane. In addition, TonB1 and FtsH are required by both pyocins to energise their transport into cells and catalyse their translocation across the inner membrane, respectively. Expression of *PA0434* was found to be specifically regulated by copper availability and we have designated PA0434 as Copper Responsive Transporter A, or CrtA. To our knowledge these are the first S-type pyocins described that utilize a TBDT that is not involved in iron uptake.

## INTRODUCTION

*P. aeruginosa* is a major cause of serious hospital acquired infections with treatment frequently complicated by high levels of antibiotic resistance with broad-resistance to β-lactams, aminoglycosides and fluoroquinolones and growing resistance to carbapenems observed globally(1). The World Health Organization (WHO) lists *P. aeruginosa* in the highest threat level of ‘critical’ for bacterial pathogens for which new antibiotics are urgently required(2).

Protein bacteriocins, such as pyocins, are narrow-spectrum antibiotics that kill bacteria closely related to the producing strain and play a key role in colonization and competition in bacterial communities(3),(4). One potential advantage of utilizing bacteriocins as therapeutic antibiotics is the ability to target a specific pathogen while avoiding collateral damage to the microbiota. This approach may enable antibiotic therapy without common complications associated with broad-spectrum antibiotics, such *Clostridium difficile* infection and domination of the microbiota with broadly drug resistant pathogens that may subsequently disseminate to cause serious systemic infection(5–7). In addition, there is growing concern that alteration of the microbiota, including that induced by the use of broad-spectrum antibiotics, may increase the risk of developing a range of chronic inflammatory diseases(8).

*P. aeruginosa* can produce a diverse range of pyocins. The soluble or S-type pyocins are multi-domain proteins produced by *P. aeruginosa* to kill strains of the same species. S-type pyocins share basic characteristics including homologous cytotoxic domains with the well-studied colicins of *E. coli*(9). Most characterized pyocins display a nuclease activity which targets DNA (S1, S2, S3, S8, Sn and AP41), rRNA (S6) or tRNA (S4 and SD2)(10,11). In addition, pyocin S5 is a pore-forming toxin and pyocin M inhibits peptidoglycan synthesis through the degradation of lipid II(12,13). Nuclease pyocins are normally co-expressed with immunity proteins that bind tightly to the cytotoxic domain to protect the producing strain from self-intoxication(14,15).

Initial contact with the cell surface for multiple S-type pyocins, including S5, SD2 and S2 and the unrelated L-type pyocin L1, is through the common polysaccharide antigen (CPA)(11,13)(16). CPA is a homopolymer of the rare deoxyhexose D-rhamnose that is not widely distributed in nature, with L-rhamnose being predominant. The CPA is therefore a useful generic receptor for diverse pyocins to target *P. aeruginosa*, enabling their accumulation on the cell surface. Structural and functional studies of pyocin S5 show that the CPA binding domain is formed by the second of two kinked three-helix bundle (kTHB) domains and that the sequence of this domain is highly conserved in pyocins that target the CPA(13). The N-terminal kTHB domains of pyocin S5 mediates interaction with the cell surface FptA transporter, a TonB-dependent transporter (TBDT). Similar to other identified pyocin transporters, the normal physiological role of FptA is in iron acquisition, in this case through transport of the iron-siderophore ferripyochelin across the outer membrane(17). Parasitisation of iron uptake pathways by pyocins extends to the energization of transport by the TonB1-ExbB-ExbD complex which services a repertoire of outer membrane TBDTs to energise outer membrane transport. In the case of pyocins, this system is hijacked through the presence of an N-terminal TonB-binding motif that directly binds to the C-terminal domain of TonB1, energizing translocation directly though the lumen of the cognate TBDT(13,18). In the case of the pore-forming pyocin S5, transport to the periplasm is sufficient for the C-terminal cytotoxic domain to insert into and depolarize the inner membrane to kill the cell. For nuclease -type pyocins, an additional inner membrane translocation domain is required to mediate transport to the cytoplasm where its nucleic acid substrate is located(19). For pyocin G, inner membrane translocation has been shown to be dependent on both the AAA^+^ATPase/protease FtsH and TonB1(20).

In this study, two novel pyocins, namely SX1 and SX2, were identified and characterized. Both pyocins display potent killing activity against *P. aeruginosa* PAO1. However, only pyocin SX2 affords effective protection in a *Galleria mellonella* model of *P. aeruginosa* infection. *In vitro* experiments demonstrated that pyocin SX1 is a metal dependent DNase. In contrast, pyocin SX2 did not show any DNase activity but inhibited protein synthesis in an *in vitro* transcription-translation assay. Mapping the uptake pathways of SX1 and SX2 shows these pyocins utilize the common polysaccharide antigen (CPA) as a cell surface receptor and a previously uncharacterized TonB-dependent transporter (TBDT) PA0434 as a translocator to cross the outer membrane. Interestingly, expression of *PA0434* was found to be regulated by copper availability hence we have designated PA0434 as Copper Responsive Transporter A, or CrtA.

## Results

### Discovery and domain organization of pyocins SX1 and SX2

To identify new pyocins we performed BLAST searches using the amino acid sequence of the pyocin S2 CPA binding and inner membrane translocation domains as query sequences(11,19). Two genes encoding putative novel pyocins, designated pyocins SX1 and SX2, were identified in the genome sequences of *P. aeruginosa* LMG5031 and *Pseudomonas sp*. 2_1_26, respectively. Both putative pyocin genes are located downstream from a P-box element which is the regulatory sequence for production of many pyocins and upstream from genes encoding putative immunity proteins(9) (**Table1**).

**Table 1.** Origin and characteristics of pyocins SX1 and SX2 and their immunity proteins.

| Strain<br>identified in | Pyocin/<br>immunity protein | Accession number | Size |  |
| --- | --- | --- | --- | --- |
|  |  |  | Amino acid | kDa |
| LMG5031 | Pyocin SX1 | KSR45004.2 | 684 | 74 |
|  | ImSX1 | KSR45005.1 | 85 | 9.9 |
| 2_1_26 | Pyocin SX2 | QKQ57980.1 | 696 | 76 |
|  | ImSX2 | QKQ57981.1 | 87 | 10 |

Comparison of the SX1 and SX2 sequences with the sequence of pyocin S2, indicates high levels of sequence identity within the CPA binding domain and inner membrane translocation domains, but little identity within the transporter binding domain (**Figure 1A**). In addition, little sequence identity between the transporter domains of pyocins SX1 or SX2 was observed with any of the characterized pyocins indicating they likely utilize a novel pyocin transporter. However, the transporter binding domains of pyocin SX1 and SX2 share 65% sequence identity indicating they may utilize the same unknown transporter. Further analysis showed that pyocin SX1 has a DNA-targeting HNH-nuclease cytotoxic domain that shares 63% amino acid identity with pyocin S2 whereas pyocin SX2 contains a pyocin G-like cytotoxic domain(19) (**Figure 1A**). A cytotoxic domain homologous to those of pyocin SX2 and pyocin G is present in carocin D and carocin S3 and from *P. carotovorum* and both of these bacteriocins have been reported to display DNase activity in vitro(21,22). The deduced domain structures for pyocin SX1 and SX2 are shown in **Figure 1B**. Based on the above analysis we predict that pyocins SX1 and SX2 utilise the CPA as their receptor and an unknown TonB-dependent receptor as their transporter.

**Figure 1.**
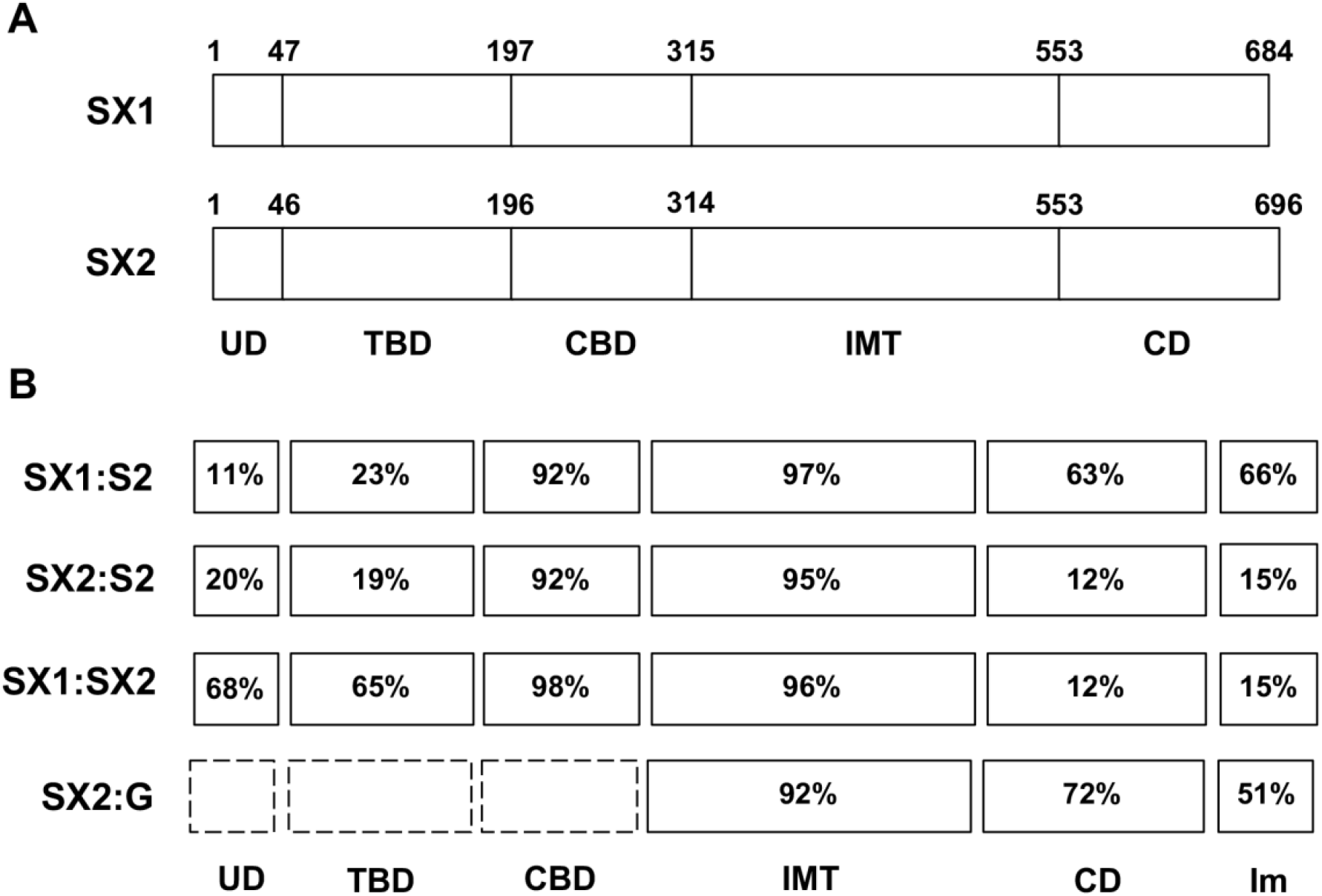
Amino acid sequence similarity of pyocins SX1 and SX2 with pyocins S2 and G and proposed domain architecture of pyocins SX1 and SX2. (A) Percentage amino acid similarity for pyocins SX1/SX2 and their immunity protein (Im) with pyocin S2 and G. (B) Proposed domain architecture of pyocins SX1 and SX2 consisting of 5 domains including N-terminal unstructured domain (UD), transporter-binding domain (TBD), CPA-binding domain (CBD), inner membrane translocation domain (IMT) and cytotoxic domain (CD). Numbers above the boxes indicate amino acid positions.

### *In vitro* and *in vivo* activity of pyocins SX1 and SX2

To determine if the putative pyocins SX1 and SX2 display killing activity against *P. aeruginosa* isolates and predicted DNase activity, we produced recombinant pyocins in complex with their cognate immunity proteins in *E. coli*. Pyocin SX1-ImSX1 and pyocin SX2-ImSX2 were isolated by Ni^2+^ affinity chromatography (via a C-terminal His_6_-tag on their respective immunity proteins) and gel filtration chromatography (**Figure 2A**). Both purified pyocins displayed potent killing activity against *P. aeruginosa* P7, using an overlay spot plate method on LB agar, with a minimum killing concentration of 0.051 µg/ml (0.6 nM) for SX1 and 0.457 µg/ml (5.3 nM) for SX2. Interestingly, pyocin SX2 also displayed efficacy in a *Galleria mellonella P. aeruginosa* infection model although pyocin SX1 did not (**Figure 2B**). Further analysis indicated that pyocin SX1 is relatively rapidly inactivated on administration to *G. mellonella* whereas SX2 retains its biological activity over an extended time period (**Figure 2C)**.

**Figure 2.**
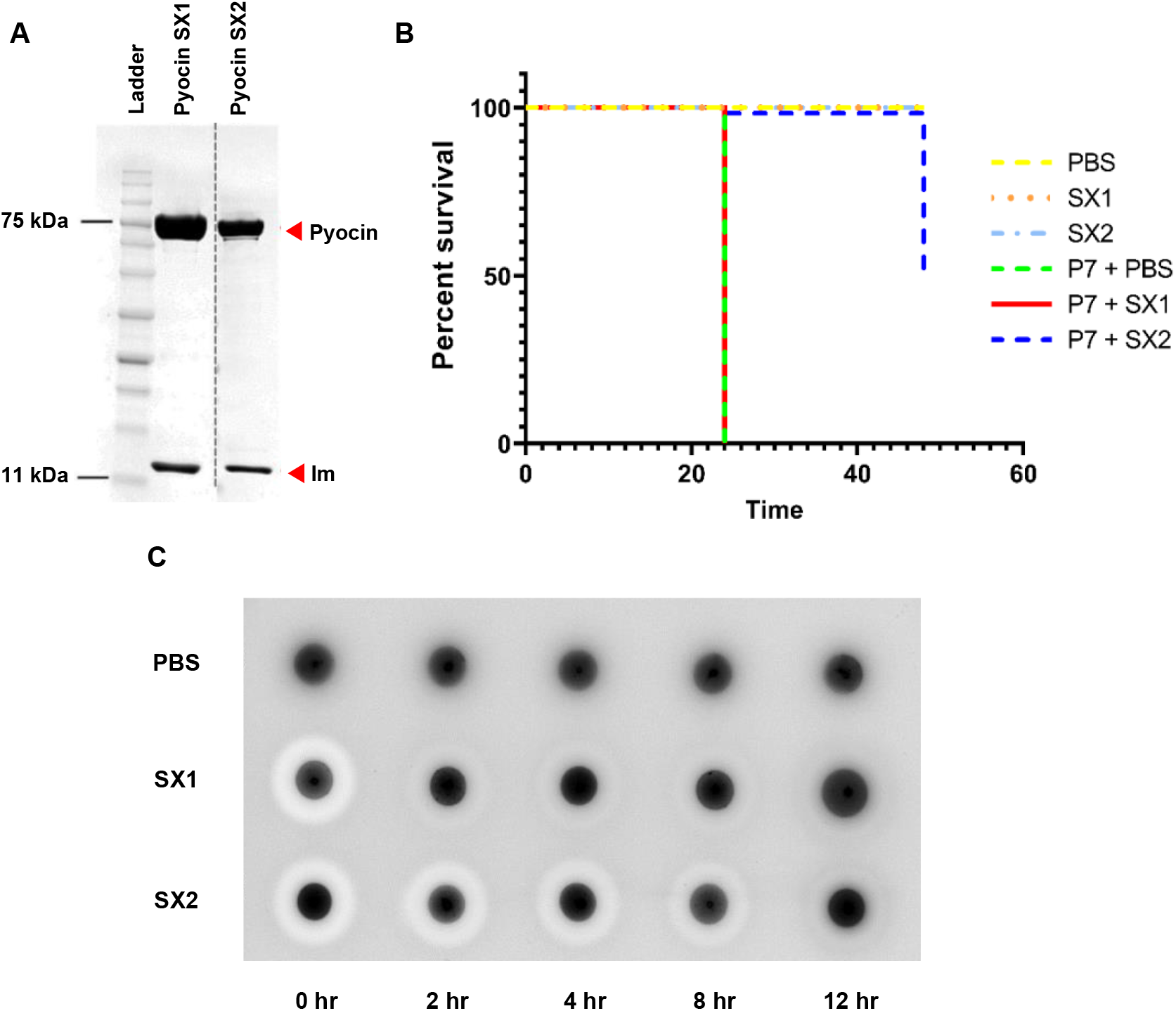
Purification and *in vivo* activity of pyocins SX1 and SX2. (A) SDS-PAGE gel (12%) of purified pyocins SX1 and SX2 (74-76 kDa) and their immunity proteins (10 kDa). The dashed line indicates splicing of gels. (B) Survival plot for groups of larvae injected with PBS or pyocins alone and groups of larvae infected with *P. aeruginosa* P7 and treated with PBS or pyocins. Groups of 30 larvae were injected with approximately 10^4^ CFU of *P. aeruginosa* P7 followed by either PBS (control), pyocin SX1 or SX2 (10 µg). The numbers of survivor were observed at 24 and 48 hr after pyocin treatment. (C) Killing activity of pyocins SX1 and SX2 after injection into *G. mellonella* larvae. Groups of 3 larvae were injected with PBS (control) or the pyocins and were collected at different time point. Three larvae were pooled, homogenized in cold PBS and centrifuged. Five microliters of the clear fraction were spotted onto *P. aeruginosa* P7 cell lawn on a LB plate.

To determine if pyocins SX1 and SX2 displayed their predicted nuclease activity against DNA, we separated the pyocins from their respective immunity proteins and tested their DNAse activity in a plasmid nicking assay. Similar to other HNH-DNase type pyocins, pyocin SX1 displayed a metal-dependent DNase activity. Pyocin SX1 was highly active in the presence of magnesium and nickel and able to completely degrade plasmid DNA, while in the presence of zinc a lower level of activity was observed (**Figure 3A**). In contrast, pyocin SX2 did not display DNase activity under any of the tested conditions (**Figure 3B**). This result is surprising given the reported DNase activity of carocin D and carocin S3, which possess a cytotoxic domain homologous to that of pyocin SX2(21,22). To probe the activity of pyocin SX2 further, we tested its ability to inhibit protein synthesis in an *in vitro* transcription-translation assay. In this experiment, 200 ng of immunity protein-free pyocin SX2, colicin D, a tRNase used as a positive control, or pyocin S5, a pore-forming pyocin, used here as a negative control, were added to the transcription-translation assay. Pyocin SX2 and colicin D significantly lowered the production of the reporter protein, Renilla luciferase, by 50% and 97%, respectively, when compared with the untreated control. The negative control, pyocin S5, had little effect on luciferase production (**Figure 3C**). In addition, the effect on luciferase production with pyocin SX2 treatment was shown to be dose dependent (**Figure 3D**). These data indicate that the cytotoxic effect of pyocin SX2 is due to interference with either transcription or translation ultimately leading to reduced protein synthesis. However, the exact molecular target of pyocin SX2 remains unknown.

**Figure 3.**
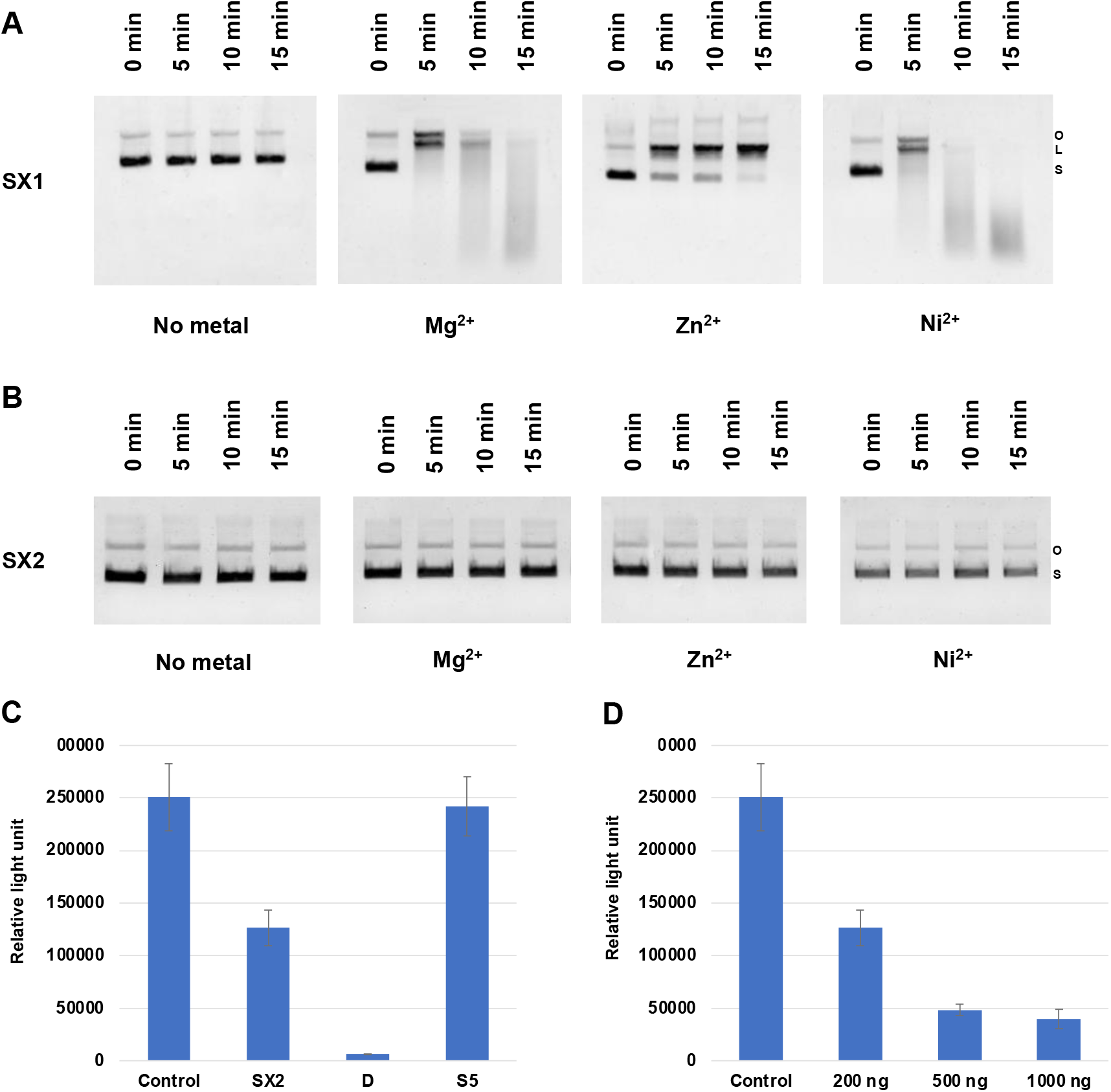
Molecular activities of pyocins SX1 and SX2. Plasmid nicking activity of pyocin SX1 and SX2. One µg of pUC18 DNA was incubated with 200 ng of pyocin SX1 (A) or SX2 (B) (immunity protein removed) in the presence different divalent metals. S = supercoiled DNA; L = linear DNA and O = open circle DNA. (C) *in vitro* transcription-translation of Renilla luciferase in the presence of 200 ng of different pyocins/colicin. (D) *in vitro* transcription-translation of Renilla luciferase in the presence of pyocin SX2 at different concentrations. The experiment was done with 3 replications. Error bars represent standard deviation of the mean. * indicate significant difference comparing to the untreated control (student’s t-test, p < 0.05).

### Identification of the receptor and transporter of pyocins SX1 and SX2

To date, all known S-type pyocins for which import mechanisms have been determined require a specific TBDT for translocation across the *P. aeruginosa* outer membrane. To identify the outer membrane transporter for pyocins SX1 and SX2, spontaneous resistant mutants were isolated by incubating late-stationary phase cells of *P. aeruginosa* PAO1 with pyocin SX1 or SX2 and plating on LB agar. The colonies grown after incubation were then picked and re-screened for pyocin sensitivity. In total, thirty spontaneous mutants isolated after pyocin SX1 or SX2 treatment were tested using spot tests for sensitivity against 4 pyocins: SX1, SX2, SD2 and L1 at 1 mg/ml. Testing against SD2 and L1 was performed to determine which isolates were likely to carry mutations that deplete or abolish CPA synthesis because both of these pyocins utilize the CPA as an outer membrane receptor(11,16). Since both SX1 and SX2 appear to carry a CPA binding domain, it was hypothesized that mutant strains resistant or tolerant to all 4 pyocins were likely to be deficient in CPA production whereas mutant strains only resistant or tolerant to SX1/SX2 were putative SX1/SX2 transporter mutants.

Consistent with this strategy, genome sequencing of 4 putative CPA production mutants showed these strains carried mutations in either the *gmd* gene encoding GDP-mannose 4,6-dehydratase (GMD) or *PA5455* coding for a putative glycosyltransferase (**Table S1**). Previous studies have shown that GMD is involved in CPA biosynthesis and *gmd* is located in the CPA O-antigen gene cluster together with other CPA biosynthesis genes such as *wzt* and *wzm*(23). *PA5455* is located in a conserved gene cluster adjacent to the CPA O-antigen gene cluster. The protein produced from this gene is predicted to be a glycosyltransferase which is proposed to be involved in CPA biosynthesis and modification(24). These data confirm that pyocins SX1 and SX2 utilise CPA as a receptor.

The genomes of 4 putative transporter mutants (resistant to both SX1 and SX2 but still sensitive to SD2 and L1) were sequenced and analyzed by comparing to the genome of the parent strain. All putative transporter mutants were found to carry nonsense mutations in the gene encoding the uncharacterized TBDT, PA0434. To confirm that PA0434 is the transporter for pyocins SX1 and SX2, both pyocins were tested against a *P. aeruginosa* strain (PAO1Δ*PA0434*) which contains a transposon insertion in *PA0434*. Both pyocins SX1 and SX2 were inactive against PAO1Δ*PA0434;* cell killing was restored after plasmid-based complementation indicating that PA0434 is the transporter for both pyocin SX1 and SX2 (**Figure 4**).

**Figure 4.**
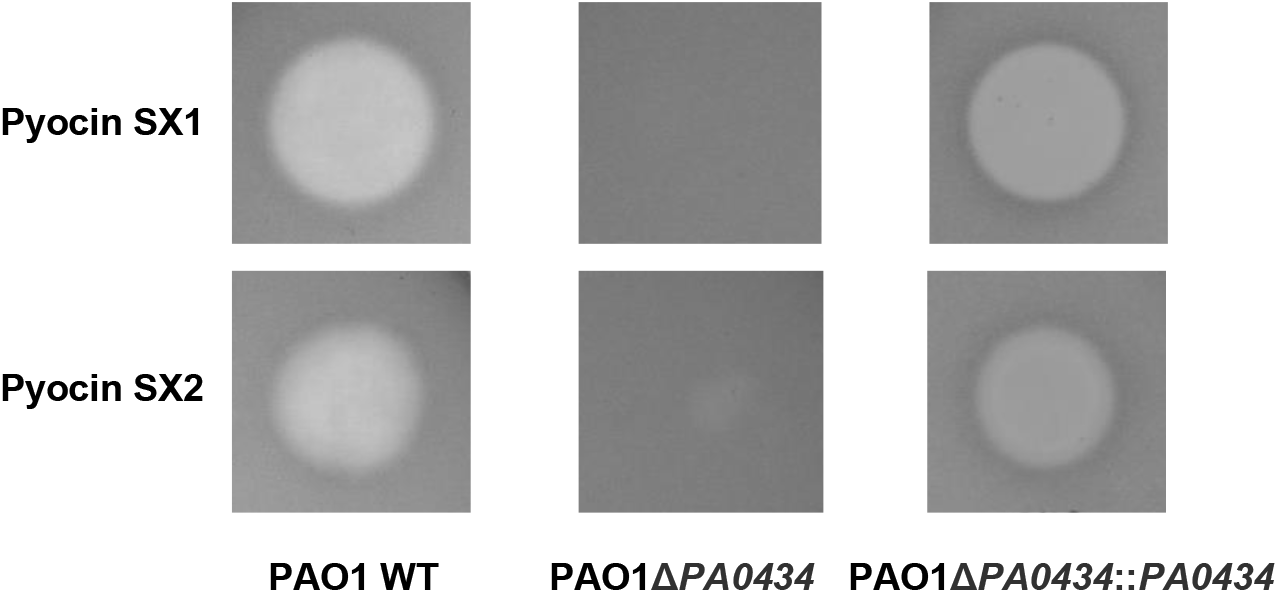
The TonB-dependent receptor PA0434 is required for pyocins SX1 and SX2 killing. Five microliters of pyocin SX1 or SX2 (1mg/ml) were spotted on to cell lawn of wild type *P. aeruginosa* PAO1 (PAO1 WT), PAO1 with transposon insertion in *PA0434* gene (PAO1Δ*PA0434*) and complement strain of PAO1Δ *PA0434* (PAO1Δ *PA0434*::*PA0434*).

Generally, the uptake of substrates via TBDTs requires an energy-transducing TonB complex, consisting of TonB, ExbB, and ExbD proteins in the inner membrane. To determine if transport of pyocins SX1 and SX2 is Ton-dependent we tested their activity against strains lacking one of the three TonB proteins encoded by the *P. aeruginosa* genome. PAO1∆*tonB1* was resistant to both pyocins SX1 and SX2 while PAO1∆*tonB2* and PAO1∆*tonB3* remained sensitive suggesting that pyocins SX1 and SX2 are TonB1 dependent (**Figure S1**). In addition, previous studies have shown that an inner-membrane protein FtsH is required for killing of nuclease-type colicins as well as pyocin G. Both pyocins SX1 and SX2 were inactive against PAO1∆*ftsH* indicating that these pyocins are also FtsH dependent (**Figure S1**).

### PA0434 is a copper responsive transporter

Killing by pyocins such as S5, SD2 and S2, which exploit iron siderophore transporters, is enhanced under iron limiting conditions, where expression of the transporter is upregulated (25). To examine the effect of Fe(III) concentration on PA0434 expression, *P. aeruginosa* PAO1 was treated with pyocins SX1, SX2 or SD2, which utilizes the ferripyoverdine transporter FpvAI, under iron-rich (with 50 µM FeCl_3_) and iron-limiting (with 200 µM 2,2’-bipyridine) conditions. Iron availability had little effect on sensitivity of PAO1 to pyocins SX1 and SX2 whereas sensitivity to pyocin SD2 was as expected inhibited by FeCl_3_ and improved by bipyridine reflecting expression of the FpvAI transporter under these conditions (**Figure S1**). These results suggest that the expression of PA0434 is independent of iron availability and so PA0434 likely does not play a role in iron uptake. Consistent with this, *PA0434* lacks an upstream Fur box (5’-GATAATGATAATCATTATC-3’) which acts as the recognition sequence for the ferric uptake regulator (Fur)(26). In *P. aeruginosa*, Fur controls both metabolism and virulence in response to iron availability including iron uptake via iron-siderophores(27). Thus, PA0434 may be involved in the uptake of a molecule other than an iron-siderophore. To determine if pyocin sensitivity can be suppressed by other metal ions, reflecting repression of PA0434 expression, the effect of Zn(II), Mn(II), Ni(II) and Cu(II) ions on sensitivity of *P. aeruginosa* PAO1 to pyocins SX1 and SX2 was determined using the overlay spot plate assay. From the metal ions tested, only the presence of CuSO_4_ decreased sensitivity of PAO1 to pyocins SX1 and SX2 while the addition of other metals did not (**Figure 5A**).

**Figure 5.**
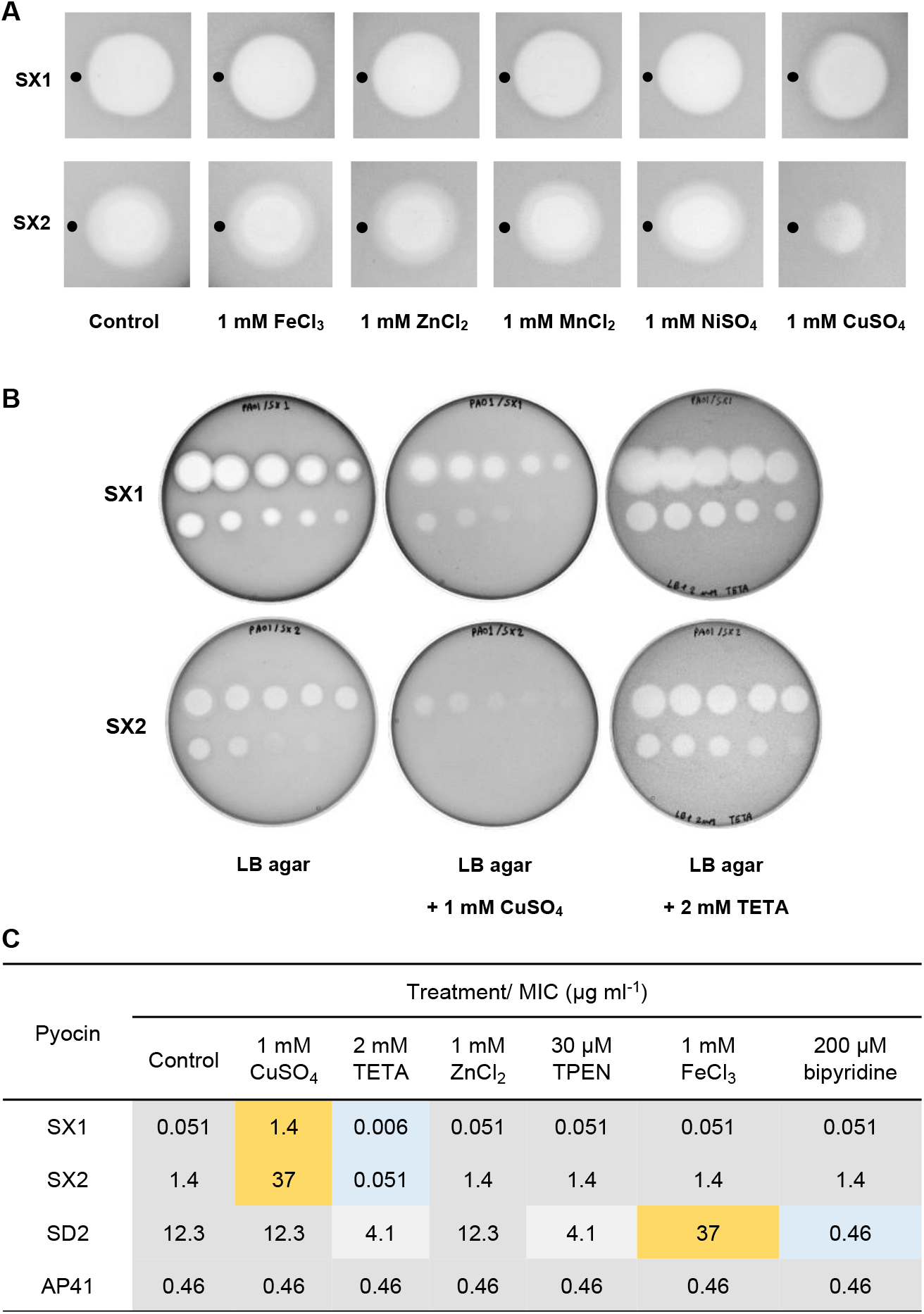
PA0434 expression specifically responds to Cu(II) availability. (A) Overlay spot plate assay of pyocins SX1/SX2 with different metal compounds spotted adjacent to the pyocin. 5 µl of 1 mM metal solution were spotted onto LB plate overlaid with *P. aeruginosa* PAO1 alongside 5 µl of 1 mg/ml pyocin SX1 or SX2 (position of metal compound spot indicated by added black circle). The overlapping areas between CuSO_4_ and pyocin spots showed decreased killing activity while other metals did not affect the killing zone indicating that Cu. (B) Overlay spot plate assay of pyocins SX1 and SX2 under normal condition (LB ager), copper-enriched conditions (LB agar + 1 mM CuSO_4_) and copper-deficient condition (LB agar + 2 µM TETA). The pyocins were serially diluted from 1000 to 0.051 µg/ml (3X dilution) and 5 µl were spotted on the plates. (C) MIC of pyocins SX1, SX2, SD2 and AP41 against *P. aeruginosa* on LB agar supplemented with different metal compounds or metal-chelators. Increasing and decreasing MICs compared to the controls are highlighted in yellow, blue and grey.

To confirm the specificity of the inhibitory effect of Cu(II), sensitivity of PAO1 to pyocins SX1 and SX2 was tested under a number of conditions. Pyocin SD2 and pyocin AP41 were selected as controls since pyocin SD2 utilizes the iron transporter FpvAI and sensitivity to pyocin AP41 is largely independent of iron availability(11,25). In these experiments, sensitivity to pyocins SX1 and SX2 was reduced when 1 mM CuSO_4_ was added to the medium. Conversely, the killing activity of both pyocins was increased in the presence of triethylenetetramine (TETA), a high-affinity Cu(II) chelator (**Figure 5BC**). FeCl_3_, ZnCl_2_, the Fe(III)-chelator bipyridine and the Zn(II)-chelator N,N,N′,N′-tetrakis(2-pyridylmethyl)ethylenediamine (TPEN) had little effect on the sensitivity of PAO1 to pyocins SX1 and SX2 suggesting that expression of the TBDT PA0434 transporter is specifically dependent on Cu(II) availability (**Figure 5C**). In addition, sensitivity to pyocin SD2 was decreased by the addition of FeCl_3_ and increased by the addition of bipyridine. The addition of CuSO_4_ and ZnCl_2_ did not affect sensitivity to pyocin SD2, reflecting the highly specific effect of iron availability on the expression of the iron-siderophore transporter FpvAI (**Figure 5C**). The killing activity of pyocin SD2 was modestly increased in the presence of TETA or TPEN. TETA and TPEN are recognized as selective copper and zinc chelators, respectively, although both chelators bind iron with a lower affinity(28,29). Sensitivity to pyocin AP41 was not affected by the presence of any metal ions or metal ion chelators (Figure 5C). The pyocin AP41 transporter has not been identified but sensitivity to the pyocin is known to be independent of iron availability(25).

To confirm that the expression of *PA0434* gene is regulated by Cu(II), *P. aeruginosa* PAO1 was grown in liquid medium with different metals or metal-chelators for 8 h and the expression level of *PA0434* gene was determined by qPCR. The expression of *PA0434* decreased by 2.5-fold in the presence of 0.5 mM CuSO_4_ and increased by 5.4-fold in the presence of 2 mM TETA. We also found that FeCl_3_ and ZnCl_2_ did not affect the expression of this gene suggesting that *PA0434* expression is specifically responsive to Cu(II) availability (**Figure 6**). A small increase in *PA0434* expression was observed in the presence of bipyridine and TPEN; however, the change in expression level was relatively small compared to TETA treatment (1.2- and 1.8-fold for bipyridine and TPEN, respectively). Expression of *fptA*, which is the gene coding for the outer-membrane transporter for Fe(III)-pyochelin and pyocin S5 was down-regulated in the presence of Fe(III) and up-regulated in the presence of bipyridine demonstrating the response of *fptA* expression to the presence of Fe(III). In addition, TETA and TPEN also triggered increased *fptA* expression, but at a lower level than bipyridine. These results indicate the effect of non-specific chelation by the metal chelators (**Figure 6**). Altogether, these results suggested that PA0434 is a Cu(II) responsive transporter and may play a role in Cu(II) transport. Thus, we named this transporter as **<u>C</u>**opper **<u>R</u>**esponsive **<u>T</u>**ransporter A or CrtA.

**Figure 6.**
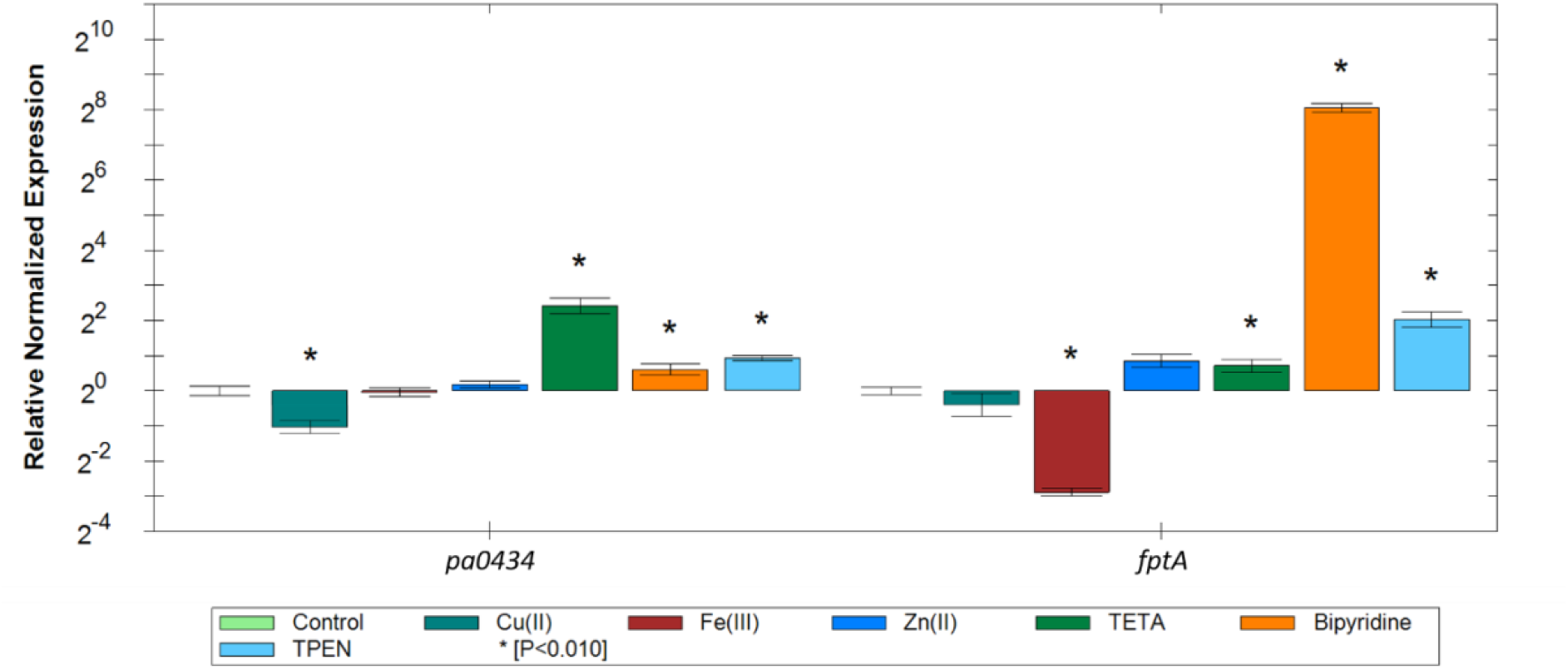
Analysis of the changes in the transcription of *pa0434* and *fptA* genes in *P. aeruginosa* cells grown with different metals or metal-chelators. RT-qPCR was performed on *P. aeruginosa* PAO1 cells grown with different metals (0.5 mM CuSO_4_, 0.5 mM FeCl_3_ or 0.5 mM ZnCl_2_) or metal-chelators (2 mM TETA, 200 µM bipyridine or 30 µM TPEN) for 8 hr. Results are given as the relative normalized expression in a Log 2 scale. The data were normalized relative to the reference gene *rpsL* and are representative of three independent experiments. Error bar indicates standard error of the mean. * indicates significantly difference compared to the untreated control (T-test, P < 0.010).

## Discussion

In this work, we describe the identification and characterization of two novel pyocins, SX1 and SX2. Pyocin SX1 was shown to be an active metal dependent nuclease that targets DNA. In contrast, pyocin SX2 did not display any DNase activity but inhibited protein synthesis in an i*n vitro* transcription-translation assay. This is a surprising result since carocin D and S3, which carry cytotoxic domains homologous to pyocin SX2 have been reported to display DNase activity. Well characterized bacteriocins, including colicin E3 are known to know kill through a targeted nuclease activity against the 16S rRNA and others, such as colicin D and E5, through cleavage of specific tRNAs. Both these activities ultimately act to inhibit protein synthesis and it remains to be determined if pyocin SX2 also targets rRNA, tRNA or acts against a different component of the RNA or protein synthesis machinery.

Both pyocin SX1 and SX2 were found to utilize CPA as a cellular receptor and a novel TBDT, CrtA (PA0434) to target *P. aeruginosa* and cross the outer membrane. Interestingly, expression of CrtA was found to be regulated by the availability of copper, suggesting that in contrast to other identified pyocin transporters, which invariably target iron-containing siderophores or heme, CrtA likely plays a role in copper uptake. Based on these results and previous studies on pyocins S2, S5 and G, a model for the transport of pyocins SX1/SX2 across the *P. aeruginosa* cell envelope is proposed (**Figure 7**). To illustrate this mechanism, the structure of the pyocin SX2-ImSX2 complex was predicted using AlphaFold-multimer(30,31). The predicted structure shows a highly elongated complex with the immunity protein bound to the cytotoxic domain, features typically associated with nuclease-type bacteriocins (**Figure 7A**). For pyocin SX2 the transporter and CPA binding domains consist of tandemly repeated kinked 3-helix bundles, as is observed in pyocin S5, and these lie N-terminal to an inner membrane translocation and cytotoxic domains. The predicted structure of the cytotoxic domain shares no similarity with other bacteriocin cytotoxic domains for which the structures are known. The mechanism we propose for pyocin SX1 and SX2 (**Figure 7B**) is a composite of the findings of this research and current knowledge derived from a variety of bacteriocins including pyocins S2, S5, G and the nuclease-type colicins(11,13,18–20,32,33). For pyocin SX1 and SX2, the translocation process is initiated by binding of the CPA binding domain to the CPA on the cell surface to localize the pyocin and allow the N-terminal unstructured domain and transporter binding domain to locate the PA0434 transporter (**Figure 7B**). Binding of pyocin SX2 to the PA0434 transporter signals for release of the TonB-box of the transporter from the membrane spanning β-barrel enabling binding of this epitope to TonB1 in the periplasm. TonB1 is able to actively induce partial unfolding of the transporter plug domain by utilization of energy from the PMF transduced by the TonB1–ExbB–ExbD complex which utilizes the proton motive allowing the N-terminal to translocate through the PA0434 barrel and present its own TonB binding epitope (TonB-box) in the periplasm. Next, TonB1 interacts with the pyocin TonB-box and mediates pyocin translocation into the periplasm. The immunity protein is released during translocation. Finally, the pyocin molecule present in periplasm is actively transported across the inner membrane via FtsH. The pyocin is likely cleaved during translocation and only the cytotoxic domain is imported into the cytoplasm, as observed for *E. coli* colicins(32,33). In addition, a recent study on pyocin G transportation across the inner membrane demonstrated that TonB1 is also required for pyocin G import into cytoplasm(19) (**Figure 7B**). TonB1 interaction may be to localize the pyocin close to the inner membrane from where the inner membrane translocation domain either interacts directly with the membrane or even with FtsH itself for transfer across the membrane and proteolytic processing(19).

**Figure 7.**
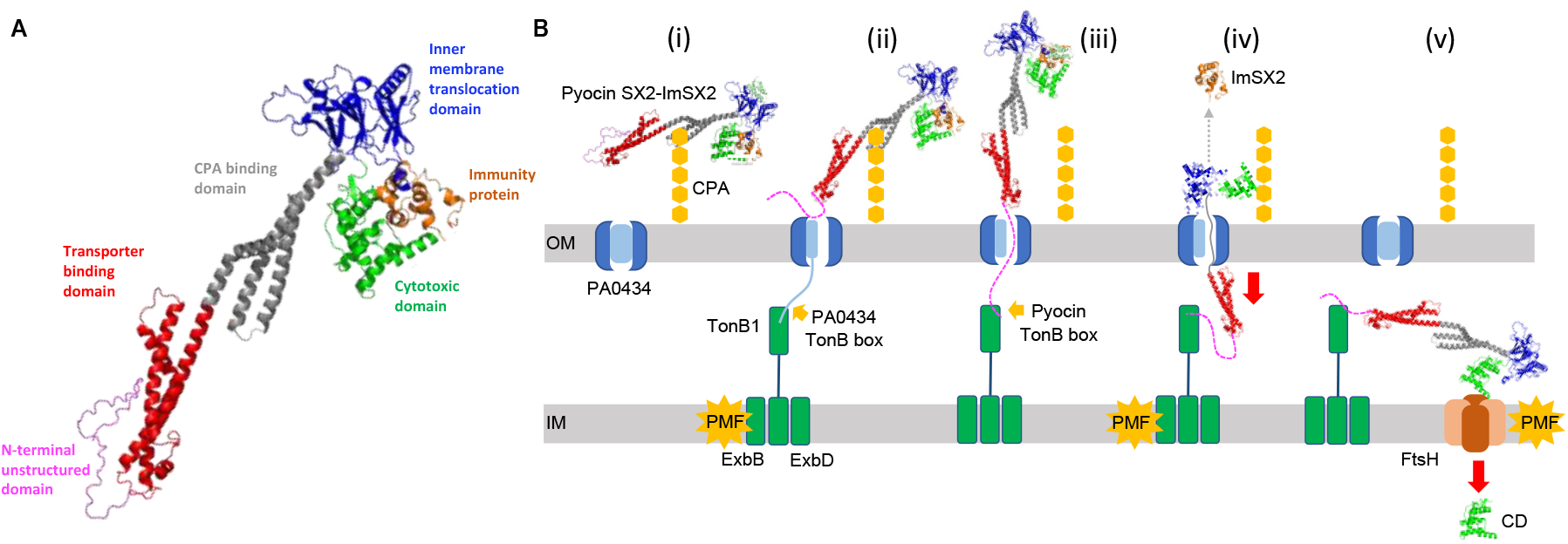
Predicted structure and entry mechanism of pyocin SX2. (A) Predicted structure of the pyocin SX2-ImSX2 complex. (B) Proposed mechanism of pyocin SX2 (and SX1) translocation into *P. aeruginosa*. (i) The pyocin CBD domain (cyan) binds to CPA on the cell surface. (ii) The UD and TBD (blue) bind to PA0434 transporter and activate unfolding of the transporter’s plug domain. (iii-iv) The UD passes through PA0434 barrel and presents its TonB-box in periplasm. Then, TonB1 binds to the pyocin TonB-box and imports the pyocin via PMF generated by TonB1-ExbB-ExbD complex. The immunity protein (Im) is released during the translocation. (v) The pyocin is proteolytically processed and only the cytotoxic domain (CD) is thought to be transported into cytoplasm via FtsH.

CrtA is a novel pyocin transporter that has not been characterized previously. Several studies have demonstrated that *P. aeruginosa* is able to utilize different siderophores produced by other microorganisms called xenosiderophores(34). A recent study on siderophore piracy showed that transcription of *crtA* gene is increased when *P. aeruginosa* was grown in metal-limited media, suggesting that CrtA likely plays a role in metal homeostasis(34). Furthermore, this study also showed the induction of *crtA* gene expression by two xenosiderophores including ferric-vibriobactin and ferric-yersiniabactin produced by *Vibrio* and *Yersinia*, respectively(34). In *P. aeruginosa*, ferric-vibriobactin is transported by a TBDT called FvbA. Even though the expression of *crtA* is induced by ferric-vibriobactin, the level of expression is relatively low when compared to the expression of *fvbA*. Moreover, *P. aeruginosa* is not able to utilize Fe(III) from ferric-yersiniabactin suggesting that ferric-vibriobactin and ferric-yersiniabactin are not the specific target for CrtA(34). However, these xenosiderophores may share some structural characteristics with the genuine target and partially trigger the expression of *crtA* gene. In addition, a study on the uptake of siderophore-drug conjugates indicated that the overexpression of CrtA in *P. aeruginosa* PAO1 increase susceptibility to several siderophore-drug conjugates including BAL030072, MC-1 and cefiderocol(35). Thus, the results obtained from this study and from previous reports suggest that the target of CrtA is likely to be a xenosiderophore involved in Cu(II) transport.

Due to the spread of MDR *P. aeruginosa*, the development of novel therapeutic approaches to treat *P. aeruginosa* infection has become essential. This work expands the repertoire of candidate pyocins for antipseudomonal therapeutic development and generates more understanding on how these proteins target and kill *P. aeruginosa*.

## Materials and Methods

### Bacterial strains and media

*P. aeruginosa* and *E. coli* isolates were grown in LB medium (10 g/L tryptone, 10 g/L NaCl, 5 g/L yeast extract (pH 7.2). All strains used in this study are summarized in **Table S2**. Strains were routinely grown at 37°C, with shaking at 180 rpm. Plasmids were propagated in *E. coli* DH5α (ThermoFisher Scientific, USA) and *E. coli* BL21 (DE3) pLysS (Agilent Technologies, USA) was used for the production of pyocins. Media were supplemented with ampicillin at 100 µg/ml (Sigma-Aldrich, USA), kanamycin at 50 µg/ml (Sigma-Aldrich, USA), chloramphenicol at 34 µg/ml (Sigma-Aldrich, USA) or gentamicin at 50 µg/ml (Gibco, USA) when required. Bacterial strains used in this study were stored in 50% glycerol at -80°C until used.

### Cloning, expression and purification of pyocin SX1 and SX2

The annotated sequences of pyocins SX1 and SX2 (Table 1) begin at the second start codon downstream form the ribosome binding size (RBS). However, most of bacterial genes start translation at the first start codon locating at 5-10 bases downstream from the RBS. Therefore, we synthesized the pyocin genes from the first start codon located at 8 bases downstream from the RBS. Genes encoding pyocin SX1 and SX2 and associated immunity proteins (GenBank records ON716475-ON716476) were codon optimized and synthesized (GenScript, USA) for expression in *E. coli* and introduced into the plasmid pET21a (+) (Merck, Germany) at NdeI/XhoI restriction sites to give pSX1ImSX1 and pSX2ImSX2, respectively. His_6_-tag were added to the C-terminal end of the immunity proteins. The plasmids were transformed into *E. coli* BL21 (DE3) pLysS (Agilent Technologies, USA) by the heat shock method. To increase the pyocin expression level, the immunity protein genes were amplified by PCR using Phusion polymerase (New England Biolabs, UK) with the following primers; 5’-ACA GAT CAT ATG GAG AAG CGT ACC ATC AGC-3’ and 5’-ATC TGT CTC GAG GCC CGC TTT AAA ACC C-3’ for ImSX1, and 5’-ACA GAT CAT ATG CTG GAC CTG GAG GG-3’ and 5’-ATC TGT CTC GAG AAC AAT ACC ATA TTT CTT TTC GG-3’ for ImSX2. The PCR products were ligated into the plasmid pET24a (+) (Merck, Germany) at NdeI/XhoI restriction sites. The plasmids with the immunity protein genes were transformed into competent *E. coli* cells carrying their counterpart pyocin genes. Expression of pyocins SX1/SX2 was induced for 4 hr by adding 1 mM IPTG into 5 L culture growth in LB broth at 37°C with shaking at 180 rpm (OD_600_ = 0.6). After induction, the cells were centrifuged, resuspended in 20 mM Tris-HCl, 500 mM NaCl and 10 mM imidazole (pH 7.5) and lysed by sonication. The cell debris was removed by centrifugation at 4°C. The cell-free lysate was applied to a 5 ml His Trap™ HP column (GE Healthcare, USA) equilibrated in the same buffer. Pyocins were eluted over a 10 - 500 mM imidazole gradient. Fractions with target pyocin were pooled, dialyzed in 50 mM Tris-HCl and 200 mM NaCl (pH 7.5), overnight at 4°C and purified by a gel filtration HiLoad 26/600 Superdex 200 pg column (GE Healthcare, USA) using the same buffer.

### Antibacterial spot assay

Eighty microliters of test strain culture (OD_600_ = 0.6) were added to 8 ml of soft agar (0.8% agar in distilled water) and poured over a LB agar plate. Five microliters of purified pyocin were spotted onto overlay plates and incubated at 37°C for 24 hr. Clear zones presented after incubation indicate killing activity.

### *Galleria mellonella* infection model

*G. mellonella* larvae were obtained from Livefood UK. *P. aeruginosa* strain P7 was grown in LB broth at 37°C until OD_600_ reach 0.6-0.7. Cells were collected by centrifugation, washed twice and diluted to OD_600_ = 0.6 in sterile PBS. To observe the protection of pyocins SX1 and SX2 against *P. aeruginosa* P7, 30 larvae were injected with 10 µl of approximately 1.5 × 10^6^ CFU/ml inoculum (100-fold dilution of OD_600_ = 0.6 bacterial suspension in PBS) into the hemocoel via the last right prolimb as previously described(36). Larvae were kept in an incubator at 37°C for 3 hr and further injected with 10 µl of pyocins at 1 mg/ml via the last left prolimb. The numbers of survivor were observed at 24 and 48 hr after pyocin treatment. For determination of pyocins remaining in larvae, larvae were injected with 10 µl of pyocins (1 mg/ml) and incubated at 37°C. At each time point, 3 larvae were transferred into a 2 ml microtube containing 400 µl of ice-cold PBS. The sample was homogenized and centrifuged at 20,000 g for 10 min at 4°C. Five microliters of clear layer were taken and spotted onto *P. aeruginosa* P7 cell lawn on a LB plate to determine pyocin killing activity.

### DNase activity assay

Immunity proteins were removed from their pyocin-immunity protein complexes by treatment with 6 M guanidine. Briefly, purified pyocin-immunity protein complex dissolved in 20mM Tris-HCl with 10 mM imidazole (pH 7.5) was applied to a 5 ml His Trap™ HP column and the pyocin was eluted by a gradient of 0 to 6 M guanidine. Pyocins without immunity protein were dialyzed in 20 mM Tris-HCl (pH 7.5) overnight before used. DNase activity assay (50 µl) was performed in 20 mM Tris-HCl (pH 7.5) with 1 µg of pUC18 DNA and one of the following metal ions, MgCl_2_ (10 mM), ZnCl_2_ (10 µM) or NiSO_4_ (10 µM). The reaction was initiated by adding 200 ng pyocin (with immunity protein removal) and incubated at 37°C for up to 15 min. The reaction was stopped by 5 mM EDTA. DNA fragments were analyzed by gel electrophoresis using 1% agarose gel.

### *In vitro* transcription and translation assay

The effect of pyocin SX2 on protein synthesis was assessed by an *in vitro* transcription-translation assay (*E. coli* T7 S30 Extract System for Circular DNA. Promega, Germany). pRL-SV40 vector (Promega, Germany) encoding Renilla luciferase was used as a template DNA. The assay was performed at 50 µl by adding 200 ng of Im-free pyocin SX2, pyocin S5 or colicin D into the reactions and incubated at 37°C for 20 min before 4 µg of the template DNA was added. The quantity of the reporter protein was examined by measuring its enzymatic activity through the Renilla Luciferase Assay System (Promega, Germany), as per manufacturer’s instruction. The light produced was measured on a Varioskan LUX microplate reader (ThermoFisher Scientific, USA). The experiment was performed with 3 replications. The tRNase colicin D and pore forming pyocin S5 were used as positive and negative controls, respectively.

### Identification of pyocins SX1 and SX2 transporter

*P. aeruginosa* PAO1 (20 independent cultures) were grown in LB broth for 48 hr to obtain late-stationary-phase cultures. One milliliter of the culture was centrifuged and re-suspended in 100 µl of pyocin solutions (1 mg/ml). The suspension was incubated at 37°C for 4 hr and spread onto a LB plate. Colonies emerging after 24 hr were selected, streaked to single colony and verified for a resistant phenotype by spot test. Genomic DNA was isolated from overnight cultures of the wild type and mutant strains using GenElute Bacterial Genomic DNA Kits (Sigma-Aldrich, USA). Library preparation and whole genome sequencing were performed by Glasgow Polyomics, UK (Illumina, paired-end, 300 bp read length and 100X coverage). CLC Genomics Workbench version 7 (Qiagen, Germany) was used for analysis of the sequences. The sequences were mapped (mismatch cost of 2, insertion cost of 3, deletion cost of 3, length fraction of 0.8, similarity fraction of 0.8) to the *P. aeruginosa* PAO1 reference genome (NC_002516.2) using CLC Map Reads to Reference tool. Subsequently, mutations were identified using by CLC basic variant detection tool with default parameters. The effects of mutations on resistant phenotype against pyocins SX1/SX2 were confirmed by using spot test on mutant strains which have transposons inserted in the genes of interest. The transposon insertion mutants were obtained from Manoil Lab, University of Washington, USA(37). A complement strain was made to confirm that a TBDT, CrtA (PA0434) is the transporter for pyocin SX1/SX2. The *PA0434* gene was amplified from *P. aeruginosa* PAO1 genomic DNA by PCR using Phusion™ High-Fidelity DNA Polymerase (ThermoFisher Scientific, USA) with the primers; 5’-ACA GAT CAT ATG AAA AAG CAC TCC ACG GCC CGC C-3’ and 5’-ATC TGT TCT AGA TCA GAA GCG CGT CTG CAC GCT CAG CTC-3’. The amplified fragment was digested with NdeI and XbaI (NEB, UK) and ligated into the modified pBBR1MCS-2 plasmid(38). The pBBR1MCS-2 plasmid was modified by introducing a NdeI restriction site at the start codon of *lacZa* gene and introducing *aacC1* gene at NsiI site. The *aacC1* gene provides resistance to gentamicin which was used as a selective marker for the compliment strain. The recombinant plasmid was transformed to *P. aeruginosa* PAO1 strain PW1793, which has transposon insertion in *PA0434* gene, by electroporation. PW1793 strain was growth overnight in LB broth at 42°C without shaking. Four milliliters of the culture were centrifuged at 14,000 g for 1 min, washed twice and re-suspended in 25 µl of 1 mM MgSO_4_. The cell suspension was mixed with 10 µg plasmid and transferred to a chilled 2 mm electroporation cuvette (Sigma-Aldrich, USA). The cells were subjected to electroporation treatment using a BioRad Gene Pulser Xcell (BioRad, USA) at 2,200 kV, 600 Ω and 25 µF. Immediately after delivery of the pulse, 975 µl of room-temperature BHI broth (Oxoid, UK) was added to the cell suspension without mixing and leaved at room-temperature for 5 min. The cell suspension was then incubated at 37°C with shaking at 180 rpm for 3 h and plated on a BHI agar supplemented with 50 µg/ml gentamicin. Colonies emerging after overnight incubation at 37°C were selected and confirmed by PCR and DNA sequencing (Eurofin, French).

### Quantitative RT-PCR

Total RNAs were extracted and purified using RNeasy Mini kit (Qiagen) together with the RNAprotect Bacteria reagent (Qiagen), as per the manufacturer’s instructions. The RNA samples were treated with DNase I (ThermoFisher Scientific, USA) and re-purified using the Monarch RNA Cleanup kit (NEB, UK). cDNA was synthesized from 1 µg of total RNA using the LunaScript RT SuperMix kit (NEB, UK) in accordance with the manufacturer’s protocol. Luna Universal qPCR Master mix (NEB, UK) was used for qPCR according to manufacturer’s recommendations (Initial denaturation 95°C, 1 min; denaturation 95°C, 15 s; extension 60°C, 30 s; 40 cycles). All reactions were prepared in a 20 µl final volume with technical duplicates and the experiments were carried out with 3 biological replicates. The primers used for RT-qPCR are as follow; 5’-TTC AGC TCG AAC CAG GTC AA-3’ and 5’-GTT GAG GAA GGA GGA CGG AC-3’ for *PA0434*, 5’-ACA TGG TGA TCA GCG GAG AA-3’ and 5’-CAG ATT CTG CTG TTC GAG GC-3’ for *fptA* and 5’-CAA GCG CAT GGT CGA CAA G-3’ and 5’-TAC ACG GCA TAC CTT ACG CA-3’ for *rpsL* gene. Absolute fold change was calculated as 2^-ΔΔct^ using *rpsL* for normalization(39,40).

## Data availability statement

GenBank records ON716475 and ON716476 contain the coding sequences for pyocin SX1 and ImSX1 from the plasmid pSX1ImSX1 and for pyocin SX2 and ImSX2 from the plasmid pSX2ImSX2, respectively.

## Supplementary Information

**Table S1.** Pyocin sensitivity and affected genes in spontaneous pyocins SX1/SX2 resistant mutants.

| Isolate | Sensitivity to pyocin |  |  |  | Affected gene | Gene product | Mutation type* |
| --- | --- | --- | --- | --- | --- | --- | --- |
|  | SX1 | SX2 | SD2 | L1 |  |  |  |
| Wild type | S | S | S | S | ND | ND | ND |
| SX1-R17 | R | R | S | S | *pa0434* | Putative TonB-dependent transporter | 1036 G del (nonsense) |
| SX1-R20 | R | R | S | S | *pa0434* | Putative TonB-dependent transporter | 68 G in (nonsense) |
| SX2-R5 | R | R | S | S | *pa0434* | Putative TonB-dependent transporter | 898 C→T (nonsense) |
| SX2-R11 | R | R | S | S | *pa0434* | Putative TonB-dependent transporter | 1764 C del (nonsense) |
| SX2-R1 | T | R | T | R | *gmd* | GDP-mannose<br>4,6-dehydratase | 450 C→G<br>(nonsense) |
| SX2-R14 | T | R | T | R | *gmd* | GDP-mannose<br>4,6-dehydratase | 379 A→T<br>(missense) |
| SX1-R6 | T | R | T | R | *gmd* | GDP-mannose<br>4,6-dehydratase | +316 G del<br>(regulation site) |
| SX2-R3 | T | R | T | T | *pa5455* | Putative<br>glycosyltransferase | 866 T del<br>868 C→A<br>(nonsense) |

The pyocin sensitivity was determined by size of clear zone produced by spot plate assay. All pyocins were tested at the concentration of 1 mg ml<sup>-1</sup>. S = sensitive; R = resistant and T = tolerant. ND indicates not detected. \* The number indicates the nucleotide position of the gene, + indicates upstream position from the gene; del = deletion; in = insertion; → = base substitution.

**Table S2.** Strains used in this study.

| Strain | Relevant characteristics | Reference or source |
| --- | --- | --- |
| ***E. coli*** |  |  |
| DH5α | *F*-, ϕ80*dlacZ*Δ*M15*,<br>Δ (*lacZYAargF*) *U169*, *deoR*, *recA1*,<br>*endA1*, *hdsR17*(*rk*-,<br>*mk*+), *phoA*, *supE44*, λ *thi*-<br>*1*, *gyrA96*, *relA1* | ThermoFisher Scientific |
| BL21(DE3)pLysS | *F*- *ompT hsdSB* (rB-mB-) *gal*<br>*dcm* (DE3) pLysS (CamR) | Agilent Technologies |
| ***P. aeruginosa*** |  |  |
| PAO1 | Clinical isolate, burn wound | (37) |
| PAO1Δ*PA0434* | Transposon insertion mutant | (37) |
| PAO1Δ*tonB1* | Transposon insertion mutant | (37) |
| PAO1Δ*tonB2* | Transposon insertion mutant | (37) |
| PAO1Δ*tonB3* | Transposon insertion mutant | (37) |
| PAO1Δ*ftsH* | Transposon insertion mutant | (37) |
| P7 | Clinical isolate, paediatric CF patient | (41) |

**Figure S1.**
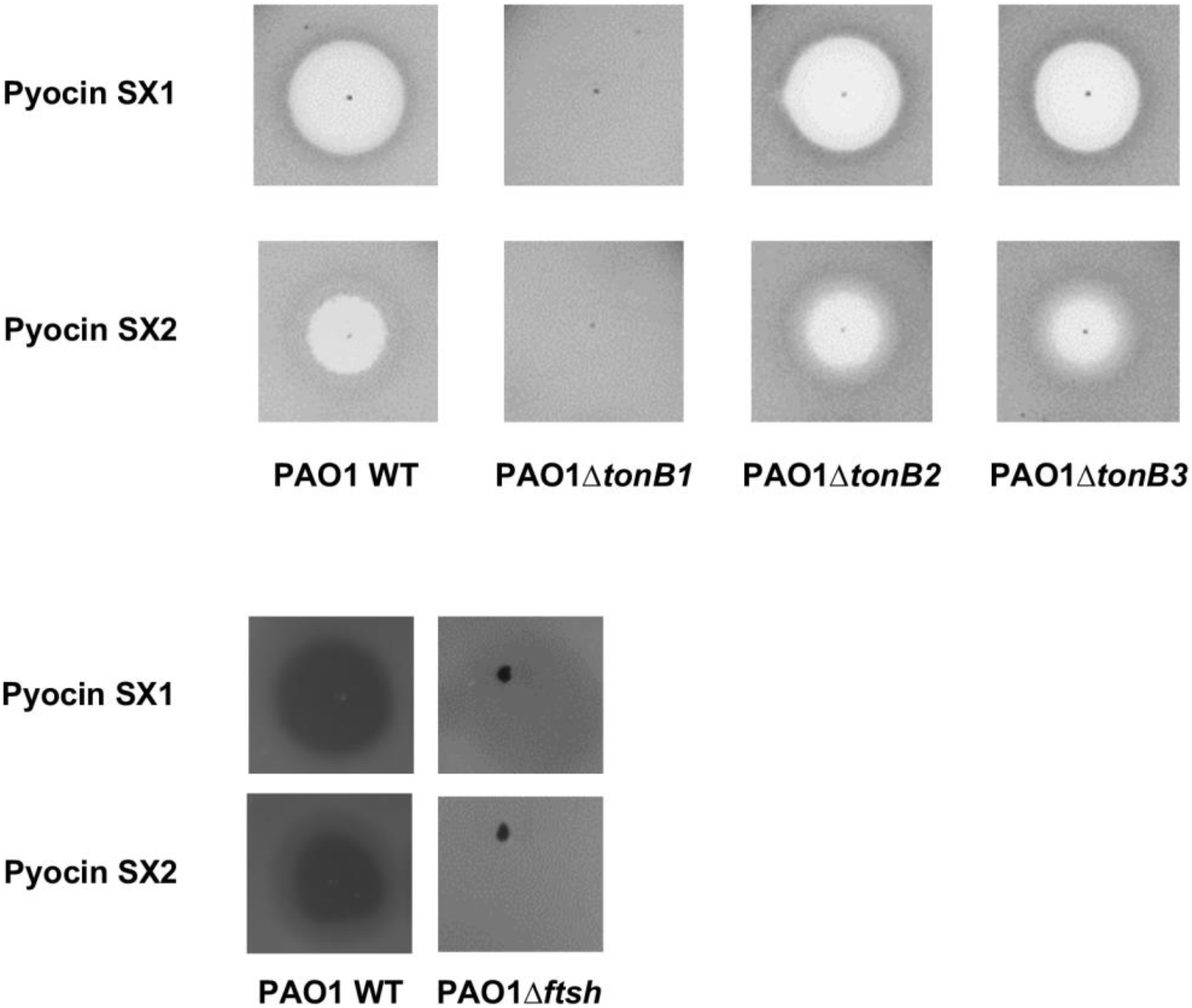
The killing activity of pyocins SX1/SX2 depend on TonB1 and FtsH. All pyocins were tested at the concentration of 1 mg ml^-1^.

**Figure S2.**
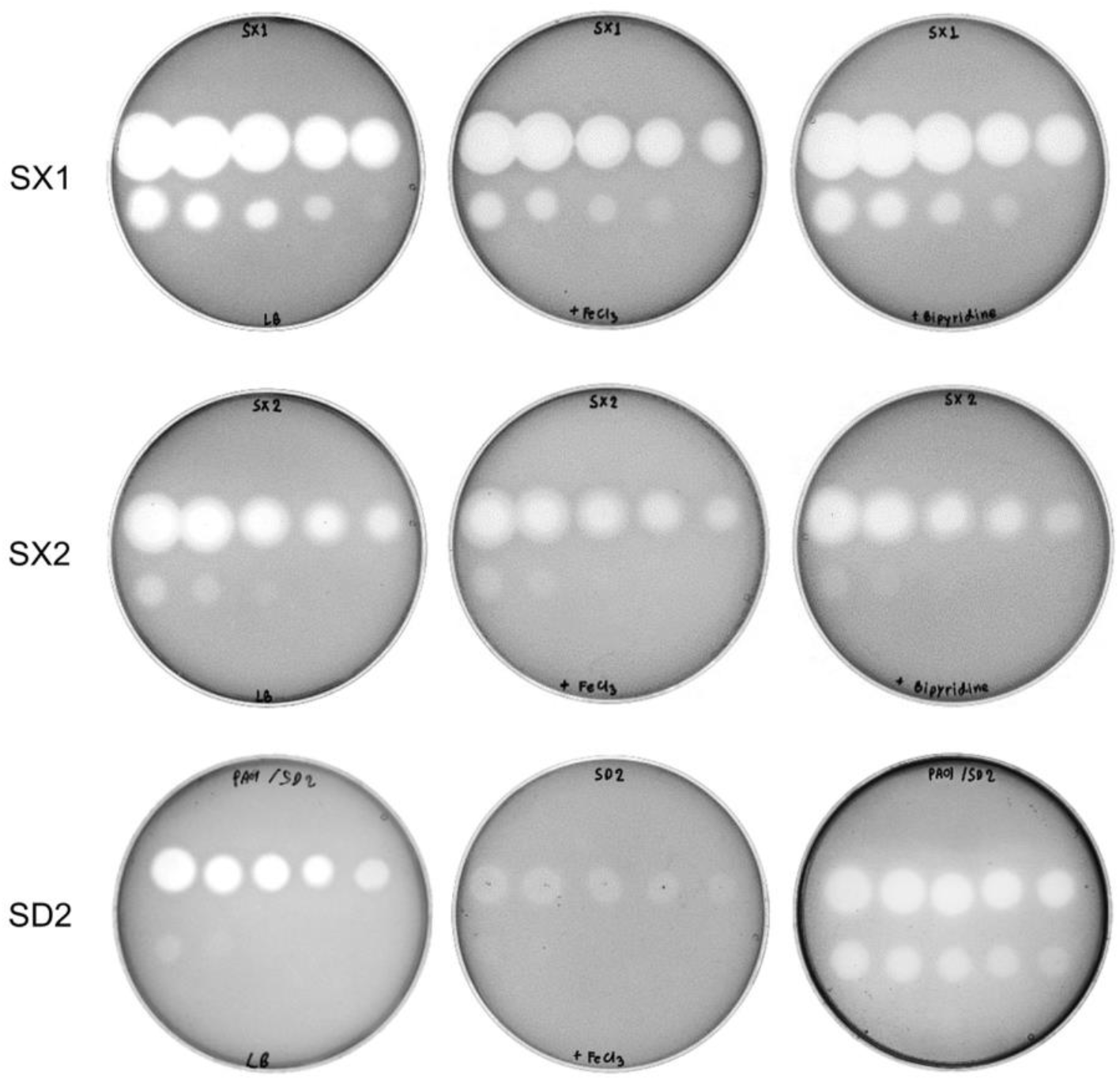
Overlay spot plate assay of pyocins SX1/SX2 and SD2 under iron-rich and iron-limited conditions. The pyocins were serially diluted from 1000 to 0.051 µg ml^-1^ (3X dilution) and 5 µl were spotted on the agar plates.

